# Characterization of Turkish Pine Honey and Differentiation from Floral Honeys by NMR Spectroscopy and Chemometric Analysis

**DOI:** 10.1101/2023.09.05.556430

**Authors:** Kerem Kahraman, Oktay Göcenler, Çağdaş Dağ

**Affiliations:** Nanofabrication and Nanocharacterization Center for Scientific and Technological Advanced Research (n^2^STAR), Koc University, İstanbul, Türkiye; Koc University Isbank Center for Infectious Diseases (KUISCID), Koc University, Istanbul, Türkiye

**Keywords:** Pine Honey, NMR Spectroscopy, authenticity, chemometric analysis

## Abstract

Honey is a viscous, supersaturated sugar solution produced by bees through the enzymatic transformation of nectar from flowers, containing a complex mixture of carbohydrates, organic acids, enzymes, and other minor constituents. Although honey has been used for thousands of years for its nutritional and medicinal properties, it has been the subject of increasing attention in recent years due to increasing adulterated honey production. Consequently, assessment of honey quality and authenticity has become essential to ensure consumer confidence of regional honey and to perverse the practice of authentic honey production. In this study, we employed nuclear magnetic resonance (NMR) spectroscopy and chemometric analysis to characterize Turkish Pine honey and compare it to flower honey originating from the Oceania, Aegean, and West Coast of North America regions. Utilizing ^1^H NMR spectroscopy, the chemical profile of Turkish Pine honey was characterized, and unique peaks were found. Additionally, PLS-DA statistical analysis was employed to further investigate the distinction of Turkish Pine honey among other floral and regional honeys. Upon completion of our statistical analysis, we were able to effectively distinguish Turkish Pine honey from other regional honey types, allowing us to formulate a universal test for authenticity of Turkish Pine Honey.

## Introduction

Honey, produced by honeybees (*Apis mellifera*), is a multifaceted product that has captivated human interest for millennia, celebrated for its nutritional, therapeutic, and cultural properties. The transformation of nectar or honeydew into honey constitutes a complex biochemical orchestration, mediated by enzymes, diastase (amylase) and invertase (α-glucosidase) (Rossano et al., 2012). Together with the intrinsic ingredients of nectar or honeydew, these enzymes give rise to a composite rich in sugars, lipids, amino acids, vitamins, minerals, and phenolic compounds (Valverde et al., 2022). It is this unique constellation of constituents that gives honey its reputed anti-inflammatory, antioxidant, antiproliferative, and antimicrobial properties (Cianciosi et al., 2018). The chemical composition of honey displays considerable variability, influenced by factors such as geographical and botanical origin, bee species, as well as yearly and seasonal conditions. Even within the same region, distinctive seasonal periods may lead to fluctuations in both the quantity and quality of honey (Tsuruda, Chakrabarti and Sagili, 2021; Walker et al., 2022). On a global scale, Turkey ranks second only to China, boasting an annual honey production capacity of 100,000 tons, encompassing diverse types such as mono-floral, multi-floral, honeydew, and pine honey (Güngör and Sen, 2018; Erdal and Tipi, 2022; Özkök, Yüksel and Sorkun, 2022; Parin et al., 2021).

Among these honey types, pine honey presents a particularly intriguing case. This unique honeydew honey, predominantly found in the East Mediterranean region (Turkey and Greece), is produced from the ripening of secretions within the honeycomb (Tananaki et al., 2007). The source of these secretions is the pine beetle, *Marchalina hellenica* (Gennadius), which deposits them on pine trees. Bees subsequently collect these secretions, infuse them with their own enzymes, and store them in beehives generating pine honey.

The medical properties and composition of pine honey have been the focus of intensive study by several research groups, uncovering a rich array of bioactive compounds (Tsavea et al., 2022; Özkök et al., 2010). Particularly, Pine honey is recognized for its antiradical effects, attributed to its phenolic compounds and amino acids. Demonstrated by Eraslan et al. (2010), the antioxidant and antiradical attributes of pine honey are well-established. Further investigations have revealed its potential antiproliferative impact, with studies highlighting the effectiveness of pine honey’s n-butanol and ethyl acetate extracts on various cancerous cell lines, including cervical, brain tumor, prostate, colon, and breast (Sıcak, Şahin-Yağlıoğlu, Öztürk, 2021).Further extending the understanding of pine honey’s composition, Duru et al. (2021) conducted a deep investigation into its volatile components leading identification of 32 distinct volatile compounds.

The authenticity of honey is intrinsically linked to its geographical and botanical origin, two vital factors that significantly influence its quality, production, and market price. As with all honey varieties, pine honey must adhere to rigorously defined quality standards before it reaches the consumer. Ensuring authenticity necessitates accurate and precise analytical methods to determine honey’s origin, thereby enhancing food quality and minimizing adulteration. Various techniques are employed in assessing honey quality, ranging from instrumental analyses for sugars and Hydroxymethylfurfural (HMF) to classical methodologies like reflectometry and titrations, and extending to more advanced techniques such as chromatographic analyses, physicochemical and sensory evaluations, vibrational and fluorescence methods, elemental and isotopic assessments, and molecular and direct MS techniques (Tsagkaris et al., 2021). Among the critical physicochemical parameters serving as authenticity markers for geographical and botanical discrimination are phenolic compounds, color, viscosity, moisture, pH, electrical conductivity, sugar content, water-insoluble matter, acidity, and HMF (Živkov Baloš et al., 2023). The interpretation of these parameters is further supported by supervised and unsupervised multivariate data analysis methods. Notably, principal component analysis (PCA) and hierarchical cluster analysis (HCA) serve as unsupervised methods, while partial least squares discriminant analysis (PLS-DA) and partial least squares regression (PLSR) function as supervised techniques (Tsagkaris et al., 2021; Walker et al., 2022).

In this context Nuclear Magnetic Resonance (NMR) spectroscopy has emerged as a particularly advantageous tool.. Its value lies in its comprehensive approach, allowing for relatively easy and rapid sample preparation and data collection. In the context of an H-NMR spectrum, the identification and elucidation of the structure of an incorrect ingredient become readily achievable when dealing with unknown compounds or impurities. NMR possesses distinct merits over alternative spectroscopic approaches, aligning with its non-destructive and non-invasive characteristics. Its environmentally friendly nature, coupled with a relatively swift and user-friendly operational paradigm, makes it suitable for routine utilization with minimal sample preparation. The technique, yielding substantial information in a concise analysis time frame, particularly excels in offering quantitative and structural insights into components within intricate mixtures, eliminating the need for pre-isolation. Additionally, the acquired data can be conveniently stored for future reference and reanalysis, enhancing the overall utility of H-NMR in scientific investigations. (Pacholczyk-Sienicka et al., 2021) Consequently, through the utilization of ^1^H NMR spectroscopy, specific product fingerprints can be determined, revealing detailed insights into the botanical origin of honey by identifying distinct organic compounds (Bertram et al., 2005). Not Surprisingly, NMR is considered as a pivotal tool in creating efficient and suitable models for honey adulteration detection (Lolli et al., 2008).

Though Turkish pine honey has been examined in various studies, a detailed analysis focusing on its specific biochemical metabolite composition and genuine characteristics is still lacking. In this study, the metabolite profiles of pine honey samples collected from Turkey were qualitatively and quantitatively characterized by NMR spectrometry. Additionally, by comparing these results with honeys from other regions, including Oceania, the Aegean, and the West Coast of North America, we aim to uncover distinctive properties of Turkish pine honey that have not been previously identified.

## 2. Material and Method

### 2.1 Sample collection

A total of thirty-three honey samples, sourced from two distinct botanical origins, including flower (n=24) and pine (n=9), were collected directly from Fijian (flower honey n=3), Hawaiian (flower honey n=6), Turkish (flower honey n=7) (Turkish pine honey n=9) and American (flower honey n=8) producers (Figure 1 and Table 1). Additionally, Manuka honey with an MGO (Methylglyoxal) level of 83 and maple syrup were bought from a local market in California. All honey samples were taken from honey produced in the year 2022. All obtained samples were subsequently stored in dark glass containers at a temperature of 4 °C prior to analysis.

**Figure 1.**
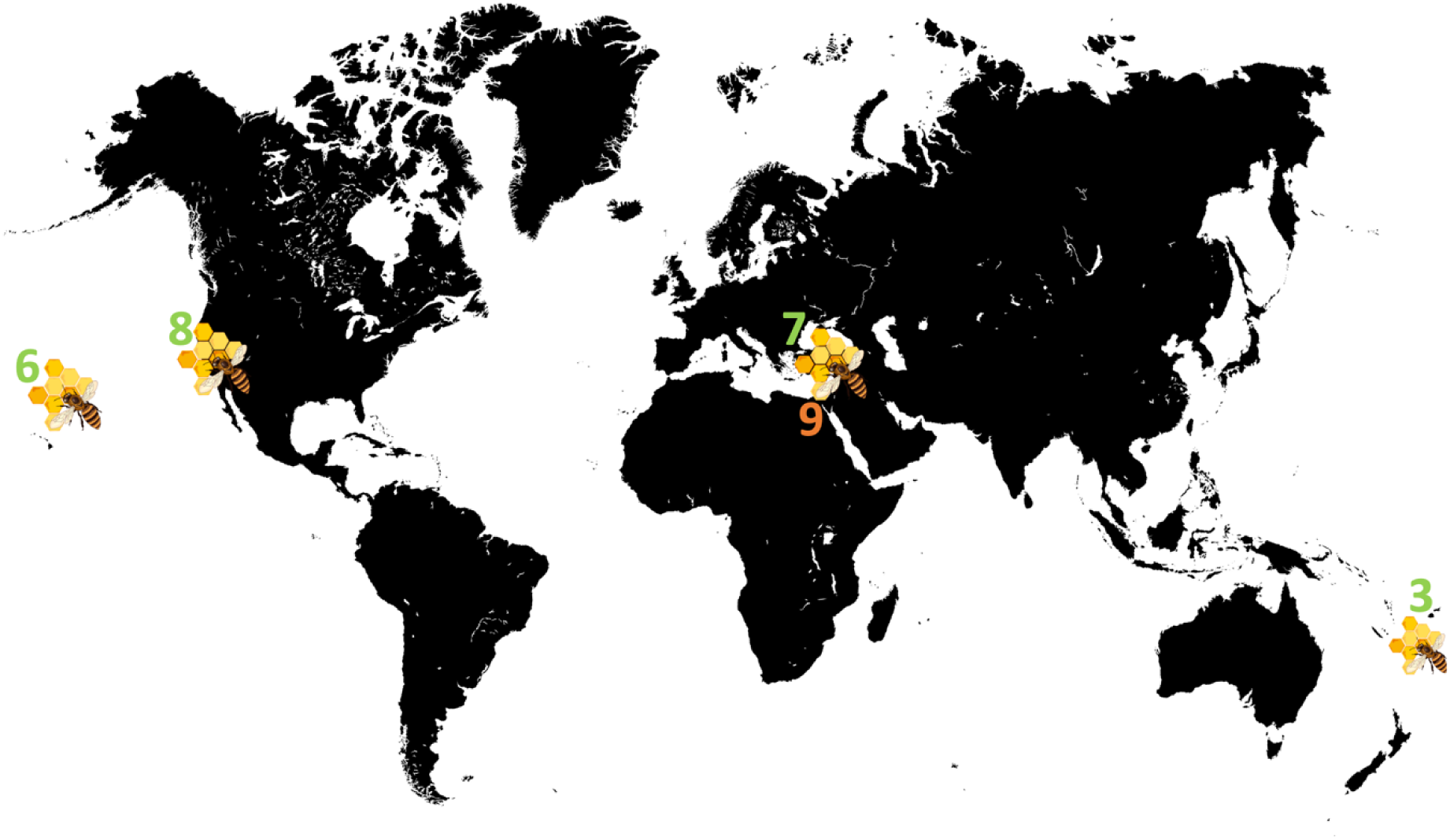
Honey Sampling Map of the study. The number of pine honey is labeled with orange and the number of flower honey is labeled with green.

**Table 1.** Sample Basic information Table.

| **Region** | **Number of Samples** | **Type of Honey** |
| --- | --- | --- |
| West Coast of America | 8 | Floral Honey |
| Fiji | 3 | Floral Honey |
| Hawaii | 6 | Floral Honey |
| western Muğla | 4 / 3 | Pine Honey / Floral Honey |
| Eastern Muğla | 5 / 4 | Pine Honey / Floral Honey |

### 2.2 Extraction and preparation of Honey samples

Honey samples were weighed in 50mL tubes and diluted 1:4 by weight using a phosphate buffer (150mM KH2PO4, pH=4.5, 100% D2O). The samples were homogenized by stirringand centrifuged at 1500 x g for 15 minutes at room temperature to remove insoluble particles. 600µL NMR samples were prepared by adding DSS (2,2-dimethyl-2-silapentane sulfonate) solution (1mM final concentration) and placed in 5mm NMR tubes.

### 2.3 Qualitative and quantitative data collection by ^1^H NMR Spectroscopy

NMR data acquisition process was completed using a 500 MHz Bruker Ascend magnet equipped with Avance NEO console and BBO double resonance room temperature probe. Water suppression experiments are commonly used for suppressing the water peak rather than routine 1D proton pulse in NMR-based metabolomics studies (Dağ et al., 2022). In this study, we used 1D NOESY-presat (noesygppr1d) pulse program for all NMR data acquisitions, and each spectrum consisted of 8K scans of 32K complex data points with a spectral width of 9615.4 Hz. Chenomx NMR Suite 9.02 software (Chenomx inc., Canada) software was used for calculation of the target metabolites (Dağ et al., 2022). Target metabolite concentrations and peak shape of other molecules calculated by Chenomx based on the concentration and peak height of the DSS. NMR data were collected with 8K scans to maximize the signal to noise ratio.

### 2.4 Statistical analysis

**MetaboAnalyst 5.0 (Pang et. al., 2021), an online software, has been used for statistical analyses. MetaboAnalyst is a user-friendly online tool that allows the creation of different types of statistical analysis results. Dataset was subjected to cube root transformation and Pareto scaling prior to analysis**. Partial least squares discriminant analysis (PLS-DA), a supervised method, was conducted for a comprehensive analysis of NMR-based metabolomics data. Partial Least Squares Discriminant Analysis (PLS-DA) represents a statistical approach primarily employed in areas like chemometrics, bioinformatics, and data analysis. This method is a specialized form of the more general Partial Least Squares regression, adapted specifically for classifying and distinguishing different groups or categories in a dataset. It focuses on pinpointing and measuring the differences that exist among various groups or classes represented in the data.

## 3. Results and Discussion

### 3.1 Analysis of Metabolite Composition in Turkish Pine Honey

Among identified water-soluble compounds in Pine Honey, 44 of them are listed with their average concentration with standard deviation in Table 2. Within the scope of the targeted metabolomics study, the NMR spectrum of each honey sample was opened and analyzed with Chenomx software. In this analysis, the DSS molecule added to all samples in equal amounts after extraction was taken into account as calibration. Thanks to its database and calibration algorithm, Chenomx software compares the peak height and volume of the DSS molecule with the peak height and shape of the target molecules and uses the calibration curves in the database to determine accurate concentrations of even metabolites at very low concentrations (Istif et al., 2023). In the analysis of Turkish Pine honey, a diverse range of chemical compounds were identified, and classified according to their chemical classifications. Carbohydrates included fructose, glucose, maltose, sucrose, arabinose, mannose, and fucose. The amino acids and related compounds included isoleucine, leucine, proline, valine, threonine, lysine, alanine, phenylalanine, aspartate, glutamine, glutamate, methylguanidine, and kynurenate. Keto acids and ketones included pyruvate, acetoacetate, acetone acetate, 2-hydroxybutyrate, lactate, acetoacetate and acetone. Ethanol was the only identified alcohol, while tartrate, fumarate, succinate, malate, citrate, 3-phenyllactate, phenylacetate, and formate were categorized as organic acids. Other classifications included, nucleotide derivatives (uridine), amides (acetamide), aromatic compounds (benzoate), other organic compounds (isocaproate, 2-hydroxyisobutyrate), quaternary ammonium cations (choline), and other nitrogen compounds (trigonelline, Pi-Methylhistidine).

**Table 2.** Identified metabolite concentrations in Turkish Pine Honey.

| Metabolite | Pine Honey<br>Average (mM) | Pine Honey<br>STD (mM) | Turkish Floral<br>Honey<br>Average (mM) | Turkish Floral<br>Honey<br>STD (mM) | Non-Turkish<br>Floral Honey<br>Average (mM) | Non-Turkish<br>Floral Honey<br>STD (mM) |
| --- | --- | --- | --- | --- | --- | --- |
| Fructose | 3875.375 | 618.867 | 19904.844 | 487.849 | 4515.246 | 1610.261 |
| Glucose | 2836.334 | 494.415 | 15978.783 | 655.755 | 3576.557 | 1062.738 |
| Maltose | 115.083 | 65.138 | 705.162 | 19.755 | 124.316 | 60.022 |
| Sucrose | 34.762 | 28.534 | 131.950 | 13.404 | 23.480 | 21.811 |
| Arabinose | 44.803 | 25.899 | 199.929 | 10.853 | 43.378 | 27.700 |
| Mannose | 7.671 | 5.438 | 38.536 | 3.836 | 4.997 | 3.680 |
| Ethanol | 1.789 | 1.328 | 8.748 | 1.225 | 4.192 | 5.560 |
| Glutamine | 2.537 | 1.230 | 9.860 | 0.613 | 2.398 | 2.071 |
| acetamide | 1.346 | 0.973 | 4.087 | 0.601 | 0.460 | 0.229 |
| lactate | 1.122 | 0.453 | 8.785 | 1.391 | 2.485 | 3.233 |
| glutamate | 4.938 | 3.751 | 16.214 | 2.516 | 4.670 | 7.326 |
| proline | 14.049 | 6.508 | 37.063 | 4.503 | 11.267 | 11.796 |
| acetate | 1.462 | 0.347 | 5.058 | 0.325 | 1.083 | 1.168 |
| threonine | 0.889 | 0.205 | 4.524 | 0.342 | 1.787 | 2.807 |
| Lysine | 3.839 | 3.193 | 11.620 | 1.629 | 1.552 | 0.703 |
| Alanine | 0.816 | 0.300 | 3.626 | 0.263 | 1.063 | 1.222 |
| Phenylalanine | 1.018 | 1.826 | 3.082 | 0.353 | 2.034 | 2.467 |
| Fucose | 0.875 | 0.271 | 3.392 | 0.266 | 12.377 | 48.389 |
| Malate | 5.646 | 4.722 | 18.339 | 2.787 | 2.690 | 1.753 |
| Choline | 1.145 | 0.354 | 4.468 | 0.373 | 1.527 | 0.906 |
| phenylacetate | 0.329 | 0.249 | 2.680 | 0.259 | 1.387 | 2.127 |
| Citrate | 2.205 | 1.858 | 4.713 | 0.586 | 1.173 | 1.433 |
| Pyruvate | 0.762 | 0.435 | 2.615 | 0.191 | 0.662 | 0.890 |
| Isoleucine | 0.425 | 0.128 | 1.508 | 0.138 | 0.691 | 0.724 |
| tartrate | 0.946 | 0.525 | 5.936 | 0.740 | 1.315 | 1.053 |
| Aspartate | 4.078 | 2.478 | 12.438 | 1.471 | 2.285 | 1.098 |
| Acetoacetate | 0.355 | 0.262 | 1.578 | 0.148 | 0.243 | 0.180 |
| tyrosine | 0.216 | 0.124 | 14.827 | 5.143 | 0.769 | 0.933 |
| Leucine | 1.627 | 3.890 | 1.210 | 0.107 | 0.652 | 0.702 |
| 3-phenyllactate | 0.337 | 0.220 | 1.398 | 0.166 | 0.995 | 1.083 |
| 2-hydroxybutyrate | 0.416 | 0.305 | 1.936 | 0.157 | 0.888 | 2.487 |
| Benzoate | 0.230 | 0.244 | 1.056 | 0.109 | 0.161 | 0.115 |
| Uridine | 0.247 | 0.133 | 1.167 | 0.090 | 0.242 | 0.194 |
| Acetone | 0.324 | 0.203 | 0.882 | 0.093 | 0.129 | 0.086 |
| Valine | 0.338 | 0.171 | 2.819 | 0.550 | 0.692 | 0.950 |
| Fumarate | 0.149 | 0.141 | 1.383 | 0.371 | 0.097 | 0.056 |
| 2-hydroxyisobutyrate | 0.168 | 0.068 | 0.524 | 0.068 | 0.129 | 0.106 |
| Isocaproate | 1.046 | 1.274 | 5.088 | 1.215 | 0.280 | 0.333 |
| Kynurenate | 0.141 | 0.115 | 0.400 | 0.028 | 0.144 | 0.098 |
| succinate | 0.409 | 0.615 | 1.320 | 0.253 | 0.148 | 0.076 |
| methylguanidine | 0.702 | 1.403 | 0.673 | 0.049 | 0.261 | 0.275 |
| formate | 0.638 | 0.826 | 1.364 | 0.394 | 1.199 | 1.007 |
| trigonelline | 0.158 | 0.051 | 0.832 | 0.048 | 0.248 | 0.131 |
| Pi-Methylhistidine | 0.302 | 0.203 | 0.862 | 0.054 | 0.449 | 0.660 |

Expectedly, the majority of compounds detected in Turkish Pine honey are readily found in numerous honeys (White, 1978). Concentrations of these compounds may differ between each honey type however their overall presence is discussed later in comparison to other honey samples. Otherwise, the presence of compounds that are not commonly found in honey acetoacetate, and Pi-Methylhistidine, are worth discussion regarding their use as potential biomarkers for authentication and adulteration detection of Turkish Pine Honey. Acetoacetate, also known as acetoacetic acid or oxobutyrate, is a short-chain keto acid that can be found in various food items, particularly various fruits and as an artificial flavoring. Soluble in water and weakly acidic in nature, it is a ketone body that also serves as an energy substrate during specific metabolic conditions such as fasting or intense exercise (Johnston et al., 1954). On the other hand, Pi-Methylhistidine is a post-translationally modified amino acid derived from the contractile proteins: actin and myosin. It is often observed in conditions associated with nitrogen loss and is excreted through urine (Elia et al., 1981). It is plausible that the appearance of these compounds in Turkish Pine honey could be a direct consequence of local organisms’ involvement in its production.

### 3.2 Development of a universal test of authenticity for Turkish Pine honey

We constructed Pairwise score plots of 5 chosen PLS-DA components to distinguish between Turkish Pine Honey and the remaining samples (Figure 2a). Following that, we selected the plot that demonstrated the two principal components responsible for the greatest degree of separation between the two groups, as shown in Figure 2b. Incorporation of a third component clearly revealed clustering of Turkish Pine honey from the rest of samples (Figure 2c). It was found that 48.5% of the total variance could be explained by the first three principal components.As depicted in Figure 2d, the VIP scores were classified into three distinct ranges, signifying the most important metabolites contributing to the separation of metabolic profiles of Turkish Pine honey from the rest of the floral honey samples. The ranges are as follows: 1.8-2.0, encompassing acetone, acetamide, and lysine; 1.5-1.8, including citrate, acetate, aspartate, and isocaproate; and 1.0-1.5, comprising tyrosine, proline, malate, 2-hydroxyisobutyrate, phenylacetate, methyl guanidine, succinate, and pyruvate. All these compounds were in higher prevalence in Pine honey, except for tyrosine and phenylacetate, which were found to be lower. Furthermore, the detailed analysis of the compounds with the most significant changes is represented by the box plots of acetamide, acetate, and acetone in Figure 2e. Among these, acetate and acetone are well-known volatile compounds found in several types of honey. Acetamide, on the other hand, deviates from this commonality, as it is not typically detected in honey.

**Figure 2.**
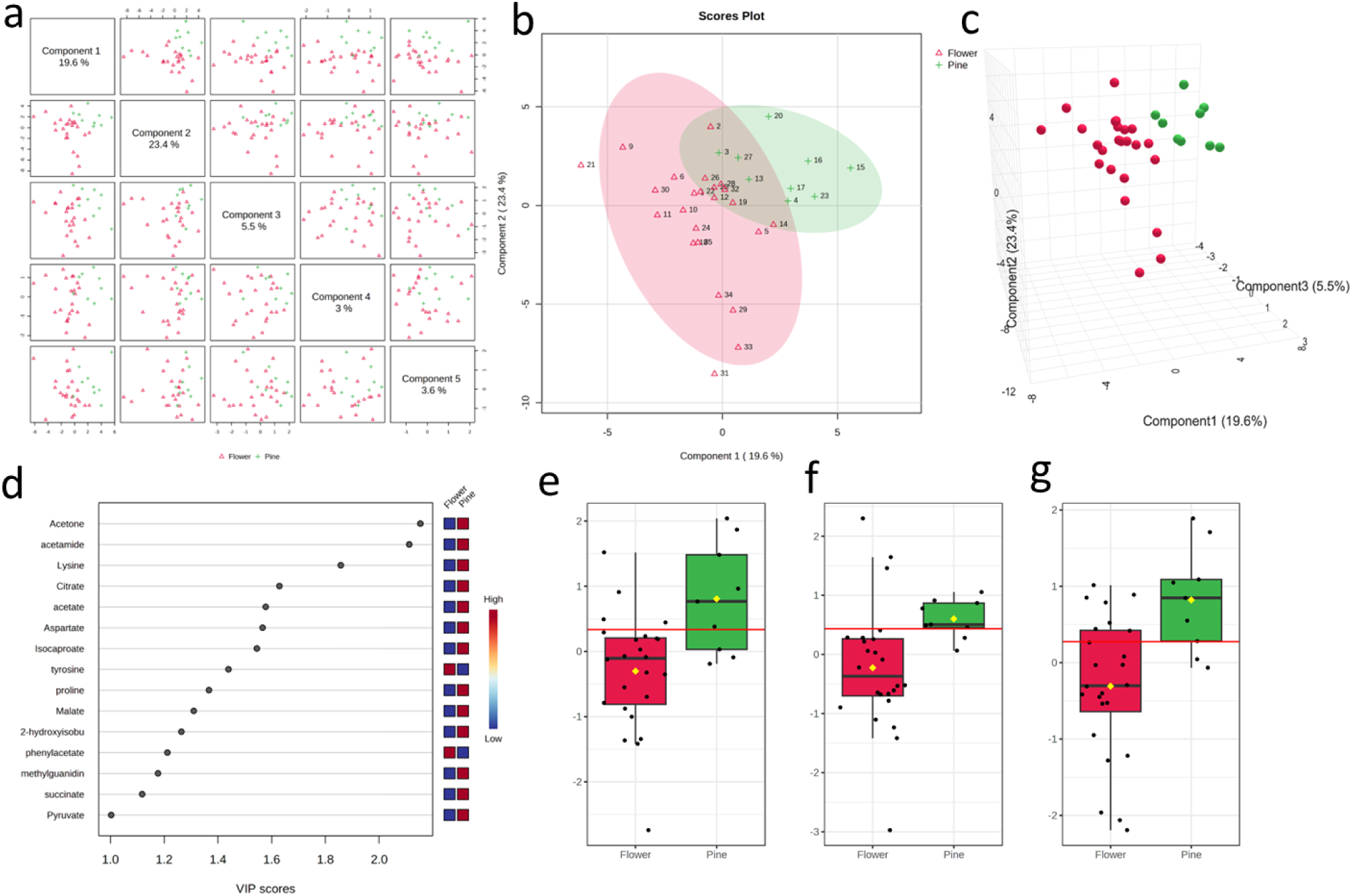
Multivariate Statistical Approach for the Discrimination of Turkish Pine Honey and Floral honey. a. PLS-DA pairwise score plot for top 5 components b. Scores plot of PLS-DA (PC1 vs. PC2) carried out on 1H NMR data of Turkish pine honeys (green) and floral honeys (red). c. 3D Scores plot of PLS-DA (PC1 vs. PC2 vs. PC3) carried out on 1H NMR data of Turkish pine honeys (green) and floral honeys (red). d. The variable importance in projection (VIP) scores obtained from the PLS-DA model. The colored boxes on the right demonstrate the relative concentrations of the corresponding metabolite. e. quantitative variation of acetamide f. quantitative variation of acetate g. quantitative variation of acetone

NMR spectrum of honey samples from different origins and maple syrup shown at figure 3a-c. The conspicuous presence of acetamide in Turkish Pine honey compared to other samples could signify a distinct chemical characteristic or metabolic pathway unique to this honey type. In addition to assigned compounds, NMR spectra were compared following peak identification (Figure 3d) to pick out peaks that are unique to pine honey samples. Peaks at 1.85, 1.87, and 1.89 ppm were observed to be unique to pine honey. These peaks can be assigned to their respective compounds and potentially be used as biomarkers to verify the authenticity of pine honey or distinguish it from other honey types.

**Figure 3.**
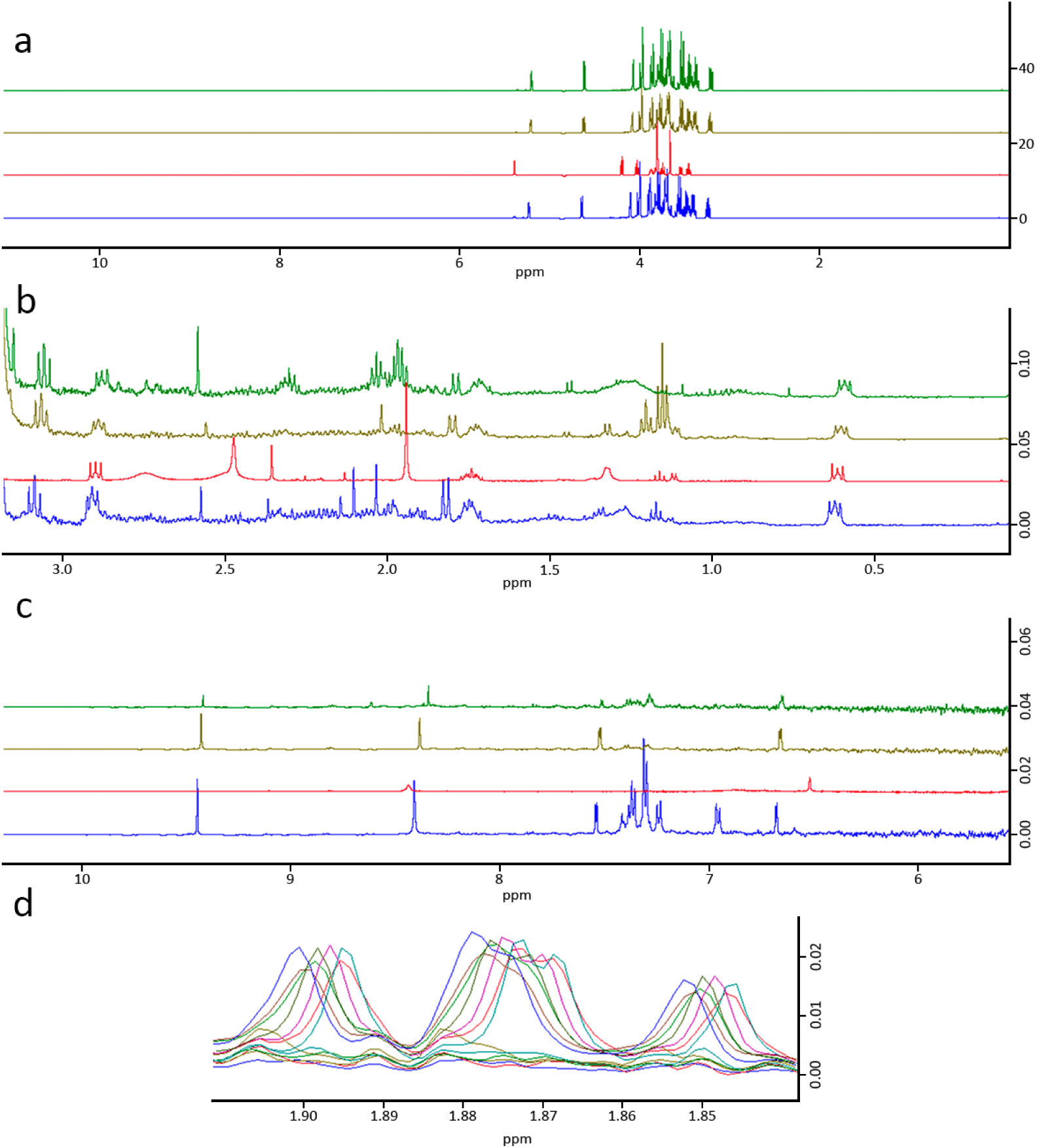
NMR spectrum of honey samples from different origins and maple syrup. a. Complete ^1^H NMR spectrum of pine honey (green), flower honey (brown), Manuka honey (Blue) and Maple syrup (red) b. 0-3.5 ppm region of ^1^H NMR spectrum of pine honey (green), flower honey (brown), Manuka honey (Blue) and Maple syrup (red) c. 5.5-10.5 ppm region of ^1^H NMR spectrum of pine honey (green), flower honey (brown), Manuka honey (Blue) and Maple syrup (red) d. Pine honey samples unique peaks at 1.89, 1.87 and 1.85 ppm (20x Zoom in to the region).

In addition to this, Figure 3 presents the general proton NMR spectra of honeys from different origins as well as maple syrup. While maple syrup can be readily distinguished at first glance due to its unique sugar content, differentiating between honey samples is not as straightforward. Notably, the peaks appearing in the 8-10 ppm range for Manuka honey are particularly striking.

We extended the analysis by constructing Pairwise score plots for the selected PLS-DA components, specifically to discern the differences between Turkish Pine Honey, Turkish Floral Honey, and Floral Honey from outside Turkey. This analysis builds upon our previous findings to further investigate the unique characteristics of Turkish Pine Honey. We selected the plot that exhibited the two principal components accounting for the most pronounced separation between these three groups, as shown in Figure 4a. This plot illustrates Turkish Flower Honey as an intermediary, bridging, characteristics of Turkish Pine Honey with Flower Honeys from outside of Turkey. As anticipated, addition of a third dimension further accentuated the clustering of Turkish Pine honey, clearly distinguishing all three groups particularly Turkish Flower Honey from the rest of the groups (Figure 4b). It was found that 44.6% of the total variance could be explained by the first three principal components. VIP scores specify the identified compounds responsible for separation mirroring the classification found in the previous analysis, however now in the context of an extended comparison (Figure 4c). In our detailed examination of the VIP scores, we identified three distinct phenomena that allowed us to discern the differential prevalence of compounds and their role in the separation. Compounds with high VIP scores over 2, such as acetone and acetamide, were found in higher prevalence in Pine Honey and, to a lesser extent, in Turkish Flower Honey and least in Non-Turkish Flower honeys. Compounds with VIP scores less than 2 such as isocaproate, malate, sucrose, and acetate also exhibited a similar pattern. Overall, these compounds can be considered as intermediary compounds which are responsible for the bridging separation of Turkish Pine Honey from Turkish Flower Honey and Non-Turkish Flower Honeys as shown in Figure 4a. Compounds, formate, lysine, glucose, trigonelline, phenylalanine, 3-phenyllactate, aspartate and citrate were found least in Turkish Flower honey on contrary to Non-Turkish Flower honey and Turkish Pine Honey. Finally, phenylacetate was also an intermediary compound between Turkish Honey and Non-Turkish Honeys with notable exception that it was found to a greater extent in Non-Turkish Flower Honeys and less in Turkish Flower Honeys and least in Turkish Pine Honey.

**Figure 4.**
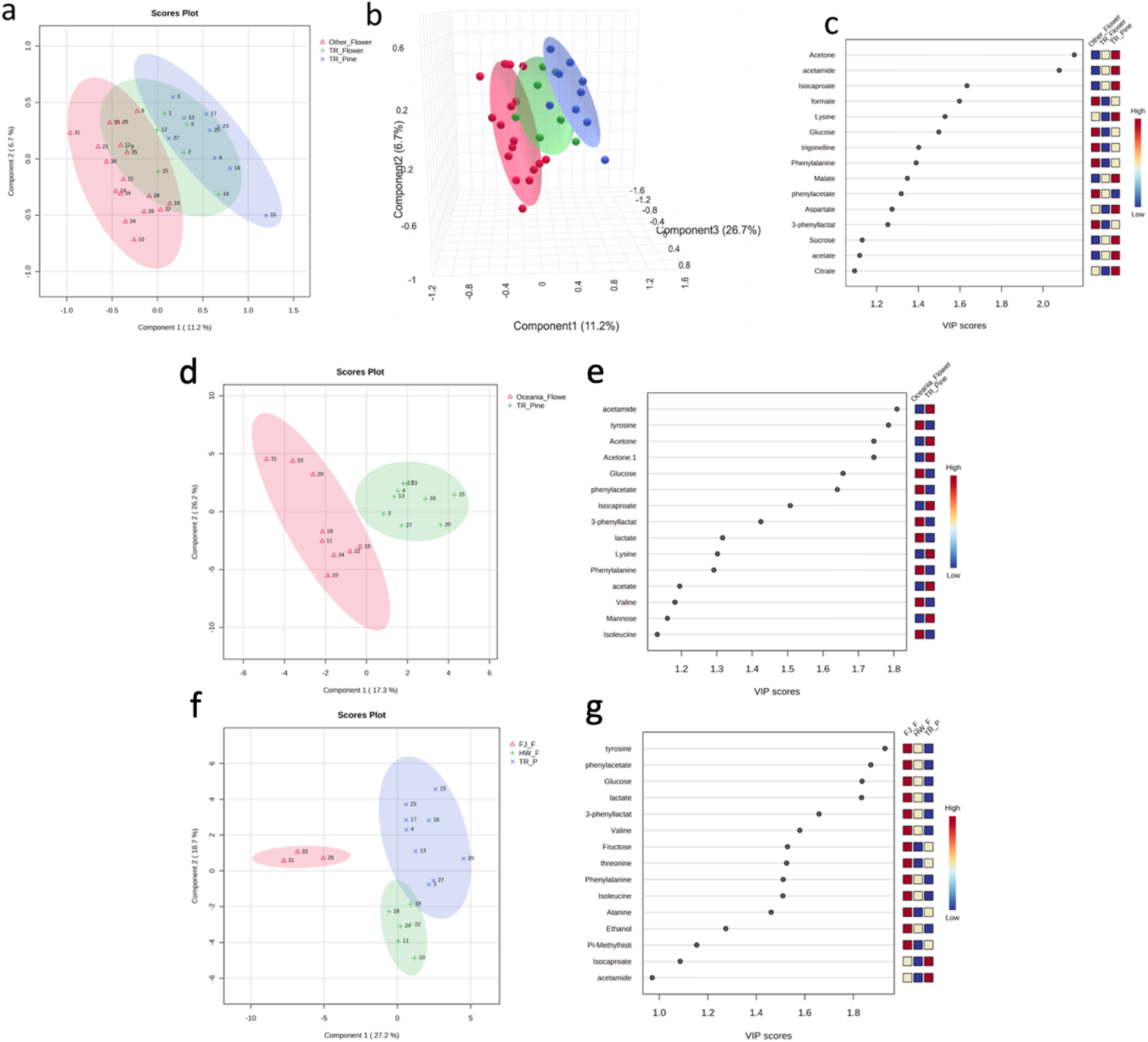
Multivariate Statistical Approach for the Discrimination of Turkish Pine Honey, Turkish Floral honey, and other Floral honey samples and specifically Oceania Floral honey samples. a. Scores plot of PLS-DA (PC1 vs. PC2) carried out on ^1^H NMR data of Turkish pine honey (blue), Turkish floral honey (green) and floral honey from out of Turkey (red). b. 3D Scores plot of PLS-DA (PC1 vs. PC2 vs. PC3) carried out on 1H NMR data of Turkish pine honey (blue), Turkish floral honey (green) and floral honey from out of Turkey (red). c. The variable importance in projection (VIP) scores obtained from the PLS-DA model. The colored boxes on the right demonstrate the relative concentrations of the corresponding metabolite. d. Scores plot of PLS-DA (PC1 vs. PC2) carried out on 1H NMR data of Turkish pine honey (green) and Oceania floral honey (red). e. The variable importance in projection (VIP) scores obtained from the PLS-DA model. The colored boxes on the right demonstrate the relative concentrations of the corresponding metabolite. f. Scores plot of PLS-DA (PC1 vs. PC2) carried out on 1H NMR data of Turkish pine honey (blue), Hawaiian floral honey (green) and Fijian floral honey (red). g. The variable importance in projection (VIP) scores obtained from the PLS-DA model. The colored boxes on the right demonstrate the relative concentrations of the corresponding metabolite

In a further effort to refine our understanding of the distinctions of Turkish Pine honey, we performed a comparative analysis between Turkish Pine Honey and Oceania Flower Honey, as well as between Turkish Pine Honey, Hawaiian Floral Honey, and Fijian Floral Honey. First, the scores plot of PLS-DA was generated to compare Turkish Pine Honey and Oceania Floral Honey, as illustrated in Figure 4d. There was clear separation between the two groups. It was found that 43.5% of the total variance could be explained by the first three principal components. The VIP scores obtained from this PLS-DA model are displayed in Figure 4e. Compounds found in higher prevalence in Turkish Pine Honey and associated with VIP scores higher than 1.5 include acetamide, acetone, and isocaproate, while those with VIP scores lower than 1.5 are lysine, acetate, and mannose. Conversely, compounds that are found to a lesser extent in Turkish Pine Honey and exhibit VIP scores higher than 1.5 are tyrosine, glucose, and phenylacetate; those with VIP scores lower than 1.5 include lactate, phenylalanine, valine, and isoleucine. Next, a score plot of PLS-DA was constructed to compare Turkish Pine Honey, Hawaiian Floral Honey, and Fijian Floral Honey, as depicted in Figure 4f. It was found that 45.9% of the total variance could be explained by the first two principal components. Though a complete separation between all three groups was observed, Hawaiian Floral Honey appeared to be more closely related to Turkish Pine Honey. The VIP scores, derived from the PLS-DA model, are presented in Figure 4g. Supporting our previous findings, compounds such as tyrosine, phenylacetate, glucose, and lactate with VIP scores higher than 1.8 were found to the greatest extent in Fijian Floral Honey, less in Hawaiian Floral Honey, and least in Turkish Pine Honey. Most likely these differences in the chemical composition are pronounced variations in climate conditions between these tropical regions and the native habitat of Turkish Pine Honey. Moreover, the inherent differences between pine honey and flower honey, stemming from the diverse floral sources and unique properties of pine nectar, further contribute to the distinct chemical profiles observed in this comparative analysis. Furthermore, all compounds except isocaproate and acetamide, which also have the lowest two VIP scores, were found to the greatest extent in Fijian Floral Honey. On the other hand, compounds such as 3-phenyllactate, valine, phenylalanine, isoleucine, and ethanol were more prevalent in Hawaiian Floral Honey compared to Turkish Pine Honey, whereas fructose, threonine, alanine, and pi-methylhistidine were less prevalent. The selection of different geographical zones, including islands and coastal areas, aimed to capture a diverse range of environmental factors influencing honey composition. The inclusion of diverse geographical zones, including islands and coastal areas, in the selection process aimed to introduce variability into the sample pool. By incorporating various honey types from different climates and regions, the study sought to simulate the broader context in which Turkish Pine honey exists among a spectrum of honey varieties. This approach was designed to provide a more comprehensive understanding of how Turkish Pine honey compares to other types across a range of environmental conditions, capturing the nuances of its chemical composition within the broader landscape of honey diversity.

### 3.3 Statistical Discrimination between Turkish Pine Honey and Turkish Floral Honey

In order to display the intrinsic metabolite differences in Turkish honey we generated pairwise score plots of the selected components as shown in Figure 5a. As formerly shown in our previous analysis there is a noticeable separation between Turkish Flower Honey and Turkish Pine Honey. VIP scores obtained from the PLS-DA analysis are depicted in Figure 5b showing most of the compounds responsible for separation except tyrosine were found to a greater extent in Turkish Pine Honey. Among these compounds, Fructose and proline were the compounds with the highest VIP scores higher than 2.0. The remaining compounds with smaller VIP scores are with respect to their VIP scores are citrate, arabinose, lysine, aspartate, glutamine, malate, methylguanidine, acetamide, formate, sucrose, acetone, and leucine.

**Figure 5.**
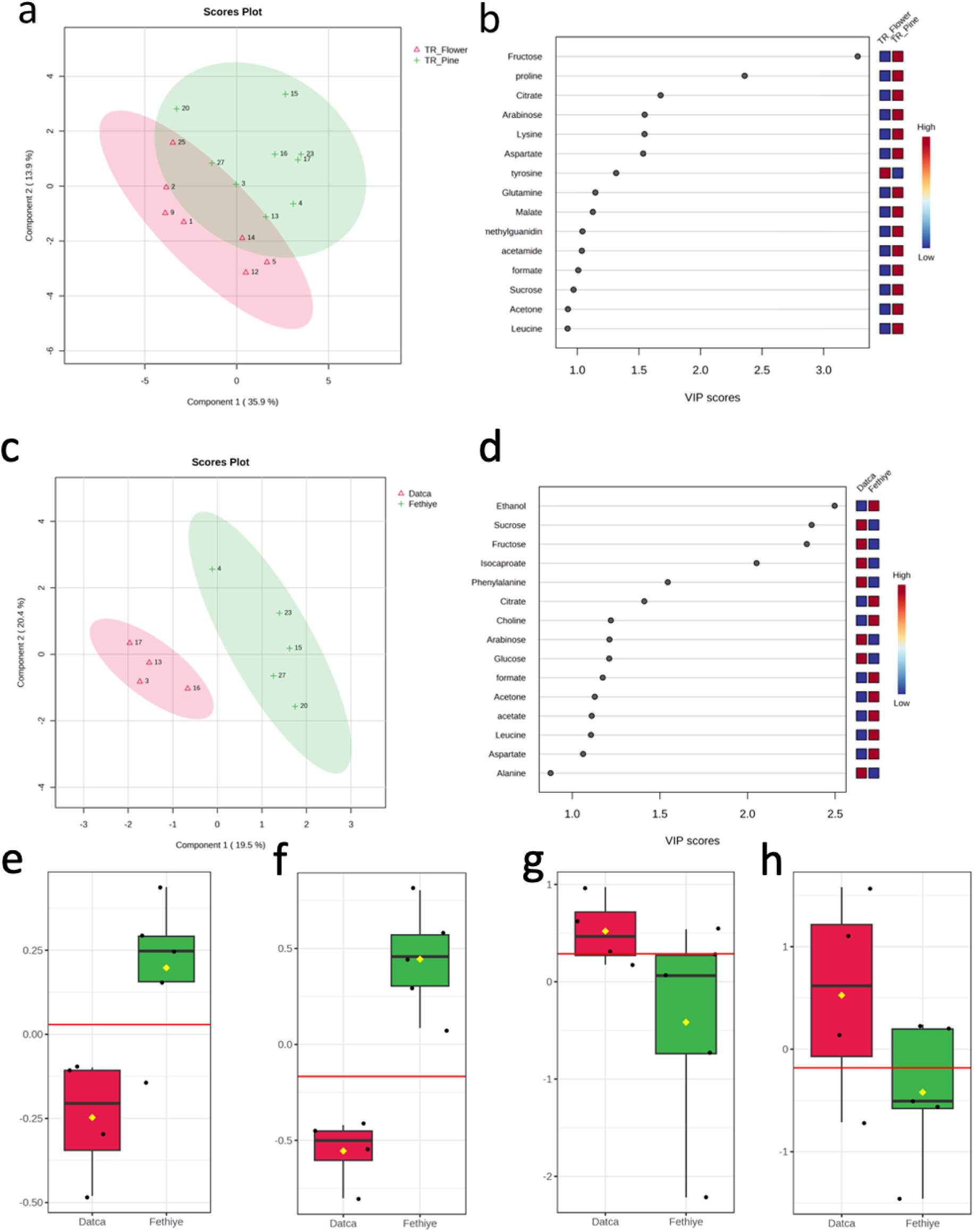
Multivariate Statistical Approach for the Discrimination of Turkish Pine Honey, and Turkish Floral honey samples and Turkish Pine Honey samples from different regions. a. Scores plot of PLS-DA (PC1 vs. PC2) carried out on 1H NMR data of Turkish floral honey (red) and Turkish pine honey (green). b. The variable importance in projection (VIP) scores obtained from the PLS-DA model. The colored boxes on the right demonstrate the relative concentrations of the corresponding metabolite. c. Scores plot of PLS-DA (PC1 vs. PC2) carried out on 1H NMR data of Turkish pine honey from western Muğla (Datça) (red) and Turkish pine honey from eastern Muğla (Fethiye) (green). d. The variable importance in projection (VIP) scores obtained from the PLS-DA model. The colored boxes on the right demonstrate the relative concentrations of the corresponding metabolite. e. quantitative variation of acetate f. quantitative variation of ethanol g. quantitative variation of fructose h. quantitative variation of sucrose

The distinctive compositional variations between Turkish Pine Honey and Turkish Flower Honey, revealed through our analysis, point toward the underlying biological and ecological factors that might be at play. Different floral sources and the unique foraging behaviors of bees may contribute to the variations in compounds such as fructose, proline, citrate, and others (Lan et al., 2021). The presence or absence of certain compounds in specific honey types could reflect the particular plants’ nectar and pollen content, soil chemistry, or the microclimate of the regions where the bees collect their materials and choice of essential floral scent compounds from fruit and vegetable crops sensory capabilities and prior experiences. (Mas et al., 2020) These intricate metabolic variations observed between Turkish Pine honey and Turkish Flower Honey offer insights into the relationships between honey composition and environmental factors. This prompts targeted exploration into the ecology and biochemistry of honey production at a specific region and time. Nevertheless such insights still have practical implications for honey authentication, quality control of commercial honey sold as Turkish Pine Honey, and potential health-related applications of Turkish honey. As demonstrated in environmental monitoring studies, honey bees and their by-products, including honey, can accumulate and reflect the presence of contaminants, thereby facilitating the detection of adulteration through deviations from established compound profiles. (Cunningham et. al., 2022)

### 3.4 Chemical composition analysis of Turkish pine honey from Mugla Province

To delve into the regional variations within Turkish Pine Honey, we conducted a comparison between samples collected from the western (Datça) and eastern (Fethiye) parts of Muğla Province. The scores plot of PLS-DA, as presented in Figure 5c, illustrates a distinguishable separation between these two regions, suggesting location-specific compositional differences. The VIP scores obtained from the PLS-DA model, depicted in Figure 5d specifying the key differences. Among compounds with VIP scores greater than 2, ethanol, which was not previously found to contribute to distinction of Turkish Pine honey, was found more prevalently in Pine honey from Fethiye whereas sucrose, fructose and isocaproate were found more in Pine honey from Datca. For compounds with lower VIP scores than 2, phenylalanine, arabinose, glucose, and alanine were found more prevalently in Pine Honey from Datca and citrate, choline, formate, acetone, acetate, leucine, and aspartate were found more in Pine honey from Fethiye. Additionally we explored the quantitative variations of specific metabolites though box plots namely acetate, ethanol, fructose, and sucrose. (Figure 5e, 5f, 5g, 5h).

The distinct separation between Turkish Pine Honey from Datça and Fethiye, as evidenced by our PLS-DA analysis, raises questions about the underlying factors influencing these compositional differences. While specific reasons for these variations cannot be definitively pinpointed without further investigation, several potential explanations warrant consideration considering these two regions are not too far apart from each other. The presence of unique floral sources in each region could contribute to differences in sugar content, such as sucrose and fructose. Ethanol, found more prevalently in Pine honey from Fethiye, might suggest variations in fermentation processes or other biogenic activities in the honey itself. Environmental factors, including climate and soil composition, might impact the availability and type of nectar collected by the bees, thus affecting the chemical profile of the honey considering Datca’s geography as a peninsula deeper into the Aegean Sea. Additionally, variations in bee foraging behaviors and hive management practices between the two locations may play a role. By deepening our understanding of regional variations in Turkish Pine Honey, we may open avenues for more targeted studies that can elucidate the factors contributing to these compositional differences. Such insights could have broader implications for the honey industry, from enhancing authenticity and traceability to optimizing production methods tailored to specific regional characteristics.

Karabagias et al. (2018) have identified and discriminated Greek Pine honey of different geographic origins using metabolite-based nuclear magnetic resonance spectroscopy (^1^H NMR) and high-performance liquid chromatography (HPLC). The obtained physicochemical data were modeled using multivariate analyses and supervised statistical methodologies.

To characterize the origin of honey, comparative volatile analysis of Greek and Turkish pine honey has been done by a purge & trap–gas chromatograph–mass (GS-MS) spectrometer system and Kohonen self-organizing maps (KSOM) algorithm (Tananaki et al., 2007). Zhang et al. (2020) have made a classification based on 1H NMR profile to separate 8 monofloral Chinese honeys, namely acacia, jujube, linden, longan, orange, rape, sunflower, vitex, into flower types and geographical origins. In the experiment, while PCA-X unsupervised modeling was carried out to determine its botanical origin, OPLS-DA supervised modeling was used to determine its geographical origin (Zhang et al., 2020). Alternating current (AC) impedance is a resistance of alternative current that can be used for evaluation of food quality and adulteration. This method is based on detecting change in impedance of a sample during stimulation which is called electrochemical impedance analysis. Hao et al. (2022) identified acacia honey adulteration by measuring the differences in AC impedance spectroscopy and metabolite analysis with 1H NMR spectroscopy. The combination of AC impedance and 1H NMR spectroscopy is a beneficial quality-control tool for identification of authenticity in honey (Hao et al., 2022).

## Conclusion

In this study, 33 honey samples were examined using 1D H-NMR spectroscopy for water soluble metabolites and subjected to PLS-DA statistical analysis to reveal differences in metabolite profiles of Turkish Pine Honey, Turkish Flower Honey, and Non-Turkish Flower Honey. Statistical analyses revealed that Pi-Methylhistidine and acetoacetate are scarcely found in most honey types and can serve as potential biomarkers for Turkish Pine Honey, as well as 1D H-NMR peaks found at 1.84, 1.87, and 1.89 ppm following assignment to their respective compounds. Acetone, acetamide, lysine, citrate, acetate, aspartate, isocaproate, proline, malate, 2-hydroxyisobutyrate, methylguanidine, succinate, and pyruvate are metabolites that were found in higher concentrations in Turkish Pine Honey samples, while Tyrosine and Phenylacetate concentrations were lower compared to floral honey types. Comparative analysis of Turkish Pine Honey, Turkish Flower Honey, and Non-Turkish Flower Honey revealed that acetone, acetamide, isocaproate, malate, sucrose, and acetate were detected in decreasing amounts for the honey types, respectively, also indicating the potential contribution of geographical factors alongside differences due to botanical origin.

PLS-DA analysis was also sufficient to observe separation between Turkish Pine Honey and Non-Turkish honey. Additionally, fructose and proline were components with the highest VIP scores when Turkish pine and floral honey samples were compared, with both being found at higher concentrations in pine honey, alongside 10 other compounds with VIP scores larger than 1, accounting for separation between the groups. Comparison between Turkish pine honey samples sourced from different locations also revealed regional differences in composition, which might be attributed to factors such as local flora, climate, and foraging behaviors of bees, although additional studies need to be conducted to address local differences.

## Author Contributions

**Çağdaş Dağ:** Conceptualization, Methodology, Supervision, Funding acquisition, Writing-Reviewing and Editing, Writing-Original draft preparation **Kerem Kahraman:** Investigation, Formal analysis, Visualization, Writing-Reviewing and Editing **Oktay Göcenler:** Investigation, Formal analysis, Writing-Original Draft, Visualization

## ACKNOWLEDGMENT

The authors acknowledge the use of the services and facilities of n^2^STAR-Koç University Nanofabrication and Nanocharacterization Center for Scientific and Technological Advanced Research. The authors gratefully acknowledge use of the services and facilities of the Koç University Is Bank Infectious Disease Center (KUIS-CID). We are grateful to Cansu Deniz Tozkoparan at Koç University for constructive feedback and for editing an earlier introduction draft of this manuscript.

## Notes

### Competing Interest Statement

The authors have declared no competing interest.

### Summary of Updates

.

## REFERENCES

Bertram, H. C., Kristensen, N. B., Malmendal, A., Nielsen, N. C., Bro, R., Andersen, H. J., & Harmon, D. L. (2005). A metabolomic investigation of splanchnic metabolism using 1H NMR spectroscopy of bovine blood plasma. Analytica Chimica Acta, 536(1–2), 1–6. 10.1016/j.aca.2004.12.070

Cianciosi, D., Forbes-Hernández, T., Afrin, S., Gasparrini, M., Reboredo-Rodriguez, P., Manna, P., Zhang, J., Bravo Lamas, L., Martínez Flórez, S., Agudo Toyos, P., Quiles, J., Giampieri, F., & Battino, M. (2018). Phenolic compounds in honey and their associated health benefits: A Review. Molecules, 23(9), 2322. 10.3390/molecules23092322

Cunningham, M. M., Tran, L., McKee, C. G., Polo, R. O., Newman, T., Lansing, L., & Guarna, & M. M. (2022). Honey bees as biomonitors of environmental contaminants, pathogens, and climate change. Ecological Indicators, 134, 108457.10.1016/j.ecolind.2021.108457

Dağ, Ç., Göçenler, O., & Tozkoparan, C.D. (2022). The analysis of metabolic content of traditional milk collected from three regions in turkey by NMR spectroscopy. GIDA, 47(5), 765–775. 10.15237/gida.GD22042

Duru, M. E., Taş, M., Çayan, F., Küçükaydın, S., & Tel-Çayan, G. (2021). Characterization of volatile compounds of Turkish pine honeys from different regions and classification with Chemometric Studies. European Food Research and Technology, 247(10), 2533–2544. 10.1007/s00217-021-03817-8

Elia, M., Carter, A., Bacon, S., Winearls, C. G., & Smith, R. (1981). Clinical usefulness of urinary 3-methylhistidine excretion in indicating muscle protein breakdown. BMJ, 282(6261), 351–354. 10.1136/bmj.282.6261.351

Eraslan, G., Kanbur, M., Silici, S., & Karabacak, M. (2010). Beneficial effect of pine honey on trichlorfon induced some biochemical alterations in mice. Ecotoxicology and environmental safety, 73(5), 1084–1091. 10.1016/j.ecoenv.2010.02.017

Erdal, B., & Tipi, T. (2022). Time Series Forecasting of Honey Production in Turkey, European Journal of Science and Technology, 35, 417–423. 10.31590/ejosat.1066665

Hao, S., Yuan, J., Cui, J., Yuan, W., Zhang, H. & Xuan, H. (2022). The rapid detection of acacia honey adulteration by alternating current impedance spectroscopy combined with 1H NMR profile. Lebensmittel-Wissenschaft + Technologie, 161, 113377. 10.1016/j.lwt.2022.113377

Istif, E., Mirzajani, H., Dağ, Ç., Mirlou, F., Ozuaciksoz, E. Y., Cakır, C., Koydemir, H. C., Yilgor, I., Yilgor, E., & Beker, L. (2023). Miniaturized wireless sensor enables real-time monitoring of food spoilage. Nature Food, 4(5), 427–436. 10.1038/s43016-023-00750-9

Johnston, J. A., Racusen, D. W., & Bonner, J. (1954). The metabolism of isoprenoid precursors in a plant system. Proceedings of the National Academy of Sciences, 40(11), 1031–1037. 10.1073/pnas.40.11.1031

Karabagias, I. K., Vlasiou, M., Kontakos, S., Drouza, C., Kontominas, M.G., & Keramidas, A.D. (2018). Geographical discrimination of pine and fir honeys using multivariate analyses of major and minor honey components identified by 1H NMR and HPLC along with physicochemical data. European Food Research and Technology, 244, 1249–1259. 10.1007/s00217-018-3040-5

Lan, J., Ding, G., Ma, W., Jiang, Y., & Huang, J. (2021). Pollen source affects development and behavioral preferences in Honey Bees. Insects, 12(2), 130. 10.3390/insects12020130

Lolli, M., Bertelli, D., Plessi, M., Sabatini, A. G., & Restani, C. (2008). Classification of Italian honeys by 2D hr-NMR. Journal of Agricultural and Food Chemistry, 56(4), 1298–1304. 10.1021/jf072763c

Mas, F., Horner, R. M., Brierley, S., Butler, R. C., & Suckling, D. M. (2020). Selection of key floral scent compounds from fruit and vegetable crops by honey bees depends on sensory capacity and experience. Journal of Insect Physiology, 121, 104002. 10.1016/j.jinsphys.2019.104002

Özkök, A., D’arcy, B., & Sorkun, K. (2010). Total phenolic acid and total flavonoid content of Turkish Pine Honeydew Honey. Journal of ApiProduct & ApiMedical Science, 2(2), 65–71. 10.3896/IBRA.4.02.2.01

Özkök, A., Yüksel, D., & Sorkun, K. (2018). Chemometric Evaluation of the Geographical Origin of Turkish Pine Honey, Food and Health, 4(4), 274–282. 10.3153/FH18027

Pacholczyk-Sienicka, B., Ciepielowski, G., & Albrecht, Ł. (2021). The application of NMR spectroscopy and chemometrics in authentication of spices. Molecules, 26(2), 382. 10.3390/molecules26020382

Pang, Z., Chong, J., Zhou, G., de Lima Morais, D. A., Chang, L., Barrette, M., Gauthier, C., Jacques, P.-É., Li, S., & Xia, J. (2021). Metaboanalyst 5.0: Narrowing the gap between raw spectra and functional insights. Nucleic Acids Research, 49(W1). 10.1093/nar/gkab382

Parin, F.N., Terzioğlu, P., Sıcak, Y., Yıldırım, K. & Öztürk, M. (2021). Pine honey–loaded electrospun poly (vinyl alcohol)/gelatin nanofibers with antioxidant properties. The Journal of The Textile Institute, 112(4), 628–635. 10.1080/00405000.2020.1773199

Rossano, R., Larocca, M., Polito, T., Perna, A. M., Padula, M. C., Martelli, G., & Riccio, P. (2012). What are the proteolytic enzymes of honey and what they do tell us? A fingerprint analysis by 2-D zymography of Unifloral Honeys. PLoS ONE, 7(11). 10.1371/journal.pone.0049164

Sen, G., & Güngör, E. (2018). Selecting Suitable Forest Areas For Honey Production Honey Production Using The AHP: A Case Study in Turkey, Cerne, 24(1).

Sıcak, Y., Şahin-Yağlıoğlu, A. & Öztürk, M. (2021). Bioactivities and phenolic constituents relationship of Muğla thyme and pine honey of Turkey with the chemometric approach. Food Measure, 15, 3694–3707. 10.1007/s11694-021-00940-8

Tananaki, C., Thrasyvoulou, A., Giraudel, J.L., & Montury, M. (2007). Determination of volatile characteristics of Greek and Turkish pine honey samples and their classification by using Kohonen self organizing maps. Food Chemistry, 101(4), 1687–1693. 10.1016/j.foodchem.2006.04.042

Tsagkaris, A. S., Koulis, G. A., Danezis, G. P., Martakos, I., Dasenaki, M., Georgiou, C. A., & Thomaidis, N. S. (2021). Honey authenticity: analytical techniques, state of the art and challenges. RSC advances, 11(19), 11273–11294. 10.1039/D1RA00069A

Tsavea, E., Vardaka, F.-P., Savvidaki, E., Kellil, A., Kanelis, D., Bucekova, M., Grigorakis, S., Godocikova, J, Gotsiou, P., Dimou, M., Loupassaki, S., Remoundou, I., Tsadila, C., Dimitriou, T.G., Majtan, J., Tananaki, C., Alissandrakis, E., & Mossialos, D. (2022). Physicochemical Characterization and Biological Properties of Pine Honey Produced across Greece. Foods, 11, 943. 10.3390/foods11070943

Tsuruda, J. M., Chakrabarti, P., & Sagili, R.R. (2021). Honey Bee Nutrition. Veterinary Clinics of North America: Food Animal Practice, 37(3), 505–519. 10.1016/j.cvfa.2021.06.006

Valverde, S., Ares, A. M., Stephen Elmore, J., & Bernal, J. (2022). Recent trends in the analysis of Honey Constituents. Food Chemistry, 387, 132920. 10.1016/j.foodchem.2022.132920

Walker, M. J., Cowen, S., Gray, K., Hancock, P., & Burns, D. T. (2022). Honey authenticity: the opacity of analytical reports - part 1 defining the problem. NPJ science of food, 6(1), 11. 10.1038/s41538-022-00126-6

White, J. W. (1978). Honey. Advances in Food Research, 24, 287–374. 10.1016/S0065-2628(08)60160-3

Zhang, J., Chen, H., Fan, C., Gao, S., Zhang, Z., & Bo, L. (2020). Classification of the botanical and geographical origins of Chinese honey based on 1H NMR profile with chemometrics. Food Research International, 137, 109714. 10.1016/j.foodres.2020.109714

Živkov Baloš, M., Popov, N., Jakšić, S., Mihaljev, Ž., Pelić, M., Ratajac, R., & Ljubojević Pelić, D. (2023). Sunflower honey—evaluation of quality and stability during storage. Foods, 12(13), 2585. 10.3390/foods12132585

